# SKEMPI 2.0: An updated benchmark of changes in protein-protein binding energy, kinetics and thermodynamics upon mutation

**DOI:** 10.1101/341735

**Authors:** Justina Jankauskaitė, Brian Jiménez-García, Justas Dapkūnas, Juan Fernández-Recio, Iain H. Moal

## Abstract

**Motivation:** Understanding the relationship between the sequence, structure, binding energy, binding kinetics and binding thermodynamics of protein-protein interactions is crucial to understanding cellular signaling, the assembly and regulation of molecular complexes, the mechanisms through which mutations lead to disease, and protein engineering.

**Results:** We present SKEMPI 2.0, a major update to our database of binding free energy changes upon mutation for structurally resolved protein-protein interactions. This version now contains manually curated binding data for 7085 mutations, an increase of 133%, including changes in kinetics for 1844 mutations, enthalpy and entropy changes for 443 mutations, and 440 mutations which abolish detectable binding.

**Availability:** The database is available at https://life.bsc.es/pid/skempi2/

## I. INTRODUCTION

Protein-protein interactions are central to almost all biological processes, from cellular signal transduction and the assembly of mesoscopic structures such as myofilaments, to viral adhesion and the immune response. Consequently the effects of changes in protein sequence on the structure, thermodynamics and kinetics of protein-protein interactions has wide implications for constraining the permissible substitutions that accrue over the course of evolution, and for understanding the molecular etiology of disease. Methods which measure, predict or optimise these changes have applications in designing *de novo* interactions [20], enhancing the specificity and affinity of biological therapeutics (e.g. [3]), designing combinatorial protein libraries (e.g. [24]), uncovering the effects of pathological mutations (e.g. [72]), locating druggable binding sites (e.g. [68]) and binding hotspots for drug design [25], altering binding kinetics [13], [64], protein-protein docking (e.g. [19]), and characterising transition states (e.g. [75]), binding pathways [61], and sequence-affinity landscapes [2].

SKEMPI is a manually curated database of mutations in structurally characterised protein-protein interactions and the effect of those mutations on binding affinity and other parameters [47]. The first release has been used as a basis for many further studies, including the development of energy functions [48], [46] which were subsequently implemented in the CCharPPI web server for characterising protein-protein interactions [49], as well as being used for ranking docked poses [45], [58], [6], [50]. SKEMPI has also been used to study human disease [56], [16], [55], assessing the role of dynamics on binding [69], exploring the conservation of binding regions [28], evaluating experimental affinity measurement methods [22], as well serving as a data source for models which predict dissociation rate changes upon mutation [1], pathological mutations [23], hotspot residues (e.g. [30], [44], [42], [66]) and changes in binding energy (e.g. [60], [17], [78], [7], [53], [51], [57], [18], [39], [76], [54], [77], [37], [5]).

Here we present a major update to the benchmark in terms of the number of mutations in the database and the number of different systems included (Table I). We now also include details of the experimental method for all entries, based on the categories of [22], as well as mutations which abolished detectable binding or for which only an upper or lower affinity limit could be ascertained for the wild-type or mutant.

**Table I.**
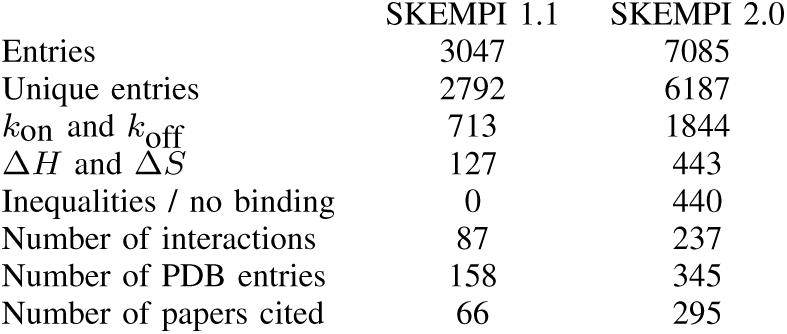
COMPARISON WITH PREVIOUS VERSION

## II. METHODS

### A. Data sources

Just over two fifths of the data come from the previous version of SKEMPI [47], comprised mostly of data found in literature sources which came to the authors’ attention, in some cases during the data collection for the structural affinity benchmark [33] and following references therein. Some entries in SKEMPI 1.1 were found by checking the references in the ASEdb [71] and PINT [36] databases, although not all the data passed the checks required for inclusion (see section II-B). Similarly, most of the new data in SKEMPI 2.0 was found by searching the literature, partly in tandem with the literature search for the more recent structural affinity benchmark [73]. During data collection three other relevant databases were published: ABbind [67], PROXiMATE [31] and dbMPIKT [41]. Their references were checked if they were not already included. Data from these sources comprise 4%, 3% and 6% of SKEMPI 2.0 respectively. As with ASEdb and PINT, none of the data were directly copied into SKEMPI. Moreover, the cited papers were read and data entered using the same checks and procedures as other entries.

### B. Data collection

Each entry was found in the literature and manually vetted. To ensure quality, a number of stringent checks were applied. Firstly, we ensured that the structure and the paper reporting the affinities refer to the same protein in the same species, and that structural and affinity data matched in terms of cofactors, ancillary chains and post-translational modifications. For instance, we distinguish between Ras^GTP^ and Ras^GppNHp^, as the nucleotide modulates the affinity of Ras with its effectors. Where the full length protein was not used, we checked to ensure that the fragment in the crystal structure matched that for which affinities are reported.

Once the checks are passed, the data is collected, including the PDB file, the chains of the interacting subunits, the mutation, the wild-type and mutant affinities (*K_D_*, *M*), the reference, the names of the proteins, the temperature at which the experiment is performed (*T*, *K*), the experimental method used (an extension of the category scheme of [22]), notes on the entry and, when available, the association rate (*k*_on_, *M*^−1^*s*^−1^), dissociation rate (*k*_off_, *s*^−1^), enthalpy (Δ*H*, kcal.mol^−1^) and entropy (Δ*S*, cal.mol^−1^.K^−1^). For cases where multiple PDB entries are available, the higher resolution structure is chosen. Where affinities or kinetic or thermodynamic parameters are reported in different units, these are converted to the units specified above. In some cases, when not reported directly, *K_D_*, *k*_on_, *k*_off_, Δ*H* and Δ*S* were calculated using the relationships Δ*G* = Δ*H* – *T*Δ*S* = *RT*ln(*K_D_*) and *K_D_* = 1/*K_A_* = *k*_off_/*k*_on_. To ensure consistency, the residue numbering in SKEMPI is the same as that reported in the PDB file. Thus the numbering is often shifted or altered compared to that in the cited paper such that, for instance, if a crystal structure of an antibody is reported in the Kabat numbering scheme but the mutation data is not, then the mutation data is converted before entry into SKEMPI. For all entries it is the case that the the affinity reported in the “wild-type” column corresponds to that of the PDB file, and the affinity in the “mutant” column is that after applying the specified mutation to the protein in the PDB file. Thus, where there are cases in which the PDB reports a mutant form and the entry corresponds to the reverse mutation back to the wild-type, the affinity of the former appears in the “wild-type” column and the latter in the “mutant” column. Such cases are noted in the database.

In addition to checking new entries, we reappraised the papers cited in SKEMPI 1.1 to collect data that were not collected previously, specifically to find mutants which abolish binding and to classify the experimental method used when not already included in the subset of SKEMPI covered in [22]. It is worth noting that often an author’s decision to report an interaction of affinity below the detection threshold as either non-binding, or as less affine than the weakest affinity presented in the paper, is arbitrary. Thus, those wishing to use the non-binding data as an inequality on the affinity may do so. We also corrected entries for 5 wild-type and 4 mutant affinities identified by [22].

### C. Post-processing and annotation

In addition to the above data, SKEMPI 2.0 also provides data on the location of the mutated residues, the homology between interactions in the data set, and processed PDB files which can be easily parsed.

#### Residue location

Each mutated residue is classified according to the scheme proposed by [38]; residues at the interface are classified as support (mostly buried when unbound and entirely buried upon binding), core (mostly solvent exposed when unbound but buried upon binding) and rim (partly buried upon binding), while residues away from the binding site are classified as interior or surface. Solvent exposed surface area was calculated using CCP4 [74].

#### Processed PDB files

The PDB files for the interactions, as downloaded from the Protein Data Bank [8], often contain multiple copies of the interacting proteins in the unit cell or other chains irrelevant to the interaction. In one instance, the binding of dimeric myostatin to follistatin-like 3, the myostatin dimer must be created by tessellating the unit cell. Further, some PDB files contain features that are not readily parsed by some software, such as residue insertion codes or negative residue numbers. To help users we provide “cleaned” PDB files which contain only the chains of interest, renumbered from one, as well as waters and other molecules with a non-hydrogen atom within 5 Å of a non-hydrogen atom of any of the chains of interest. Consequently each mutation is reported with both PDB numbering and renumbered.

#### Defining homologous interactions

Each entry also specifies which other entries are mutations to homologous interactions. Two interactions are deemed homologous if they have a shared binding partner or homologous binding partner and at least 70% of the corresponding interface residues are common to both interactions. We determine the homology between proteins using the GAP4 program [29], and define homologous proteins as those with a similarity score greater than 50 and at least 30% sequence identity. Interface residues are defined as those with a non-hydrogen atom within 10 Å of a non-hydrogen atom on the binding partner. Interactions falling within manually assigned clusters of homologous interactions are designated as pMHC/TCR, antibody/antigen or protease/inhibitor. While the names of these clusters have been chosen to reflect the predominant function of their constituent interactions, they reflect the homologies within the data set and are not functional assignments. Thus, for instance, some nanobodies are classified as antibodies as they bind to the same site as cetuximab, 14.3.d is classified as TCR, even though it is only the *β* chain, and its binding partner, enterotoxin C3, is classified as a pMHC.

## III. RESULTS & DISCUSSION

### A. Diversity, bias and interrelationships within SKEMPI

In total 7085 entries were collected, summarised in Table I and Figure 1A. These data were derived from the literature and consequently, while encompassing a broad range of residues, proteins, interactions and systems, are biased toward the interests and capabilities of the research community. These biases are evident in the composition of the database according to parameters shown in Figure 1. The ΔΔ*G* values span a large range, but mostly fall within −3 to 7 kcal.mol^−1^ (Figure 1B), for both biophysical and technical reasons (see Section III-C). Almost three quarters of the data correspond to single point mutations, and more than half of those are mutations to alanine (Figure 1C). Charge swap mutations and mutations between aromatic residues are also over-represented. Most single point mutations are located at the binding site, and most of those are at the core of the interface. Similarly, most double mutations are both in the binding site, and most of those are both in the core. By far the most popular methods for measuring binding affinity were surface plasmon resonance and spectroscopic methods such as fluorescence. Large biases towards specific interactions and classes of interaction are also present, such as early studies into protease inhibition and immunological interactions such as antibody-antigen complexes, the recognition of peptides presented on cell surfaces by T-cell receptors, cytokine signalling and the complement system. Indeed, almost half of the data corresponds to protease-inhibitor, antibody-antigen and pMHC/TCR interactions alone. While many interactions within these classes share common binding sites or homologous binding sites (Figure 2), there are also connections between these groups, for instance via inhibitory antibodies which bind to a protease active site, or due to common binding regions of proteins in the immuoglobulin superfamily, such as antibodies, TCRs, MHCs and *β*-2 microglobulin. Also present in the data are smaller clusters of shared and homologous interactions, such as the Ras-effector cluster. These relationships are noted in the database and may be useful for avoiding overfitting when developing models or for validation and estimating generalisation error, as described previously [47].

**Fig. 1.**
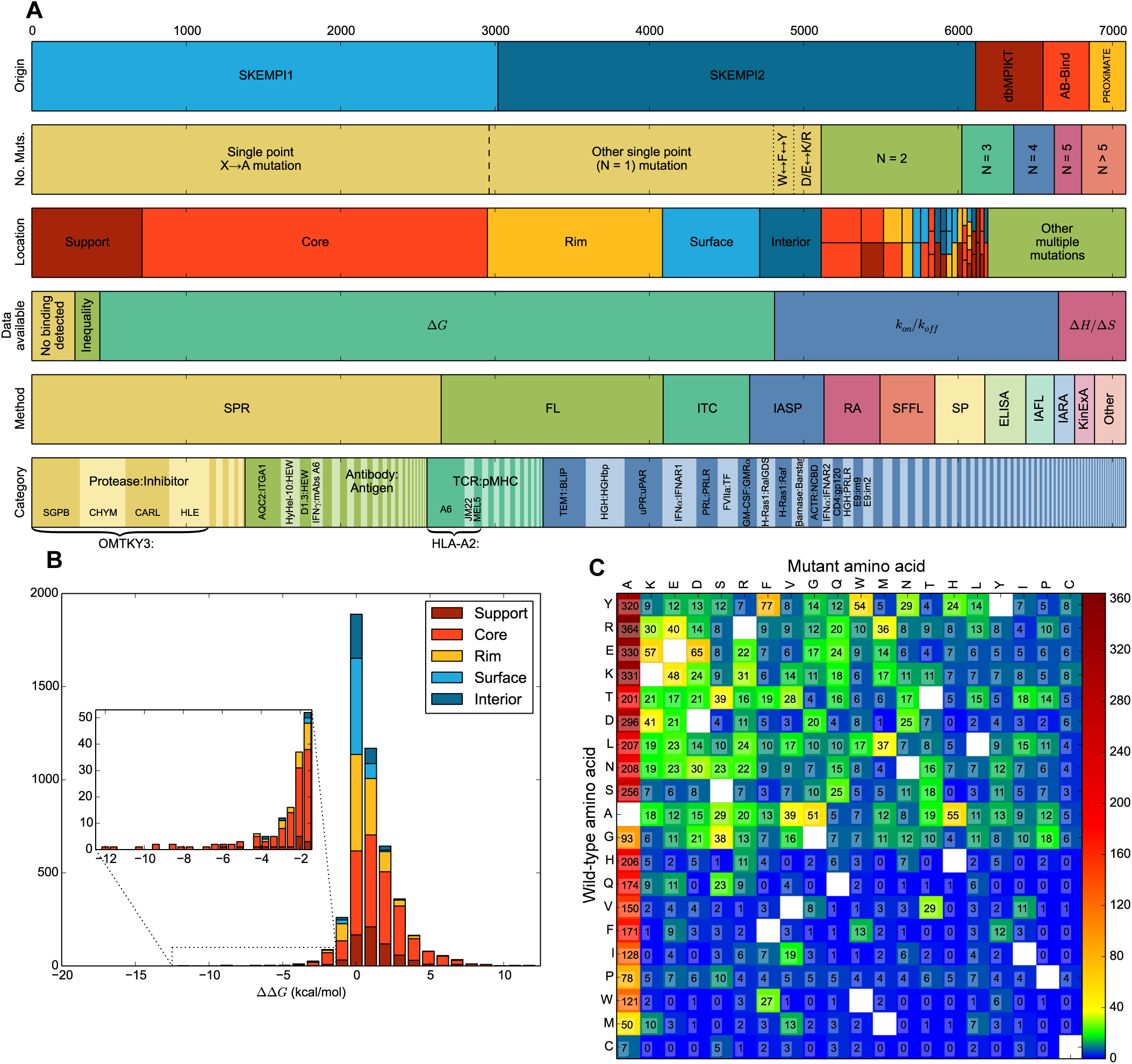
An overview of SKEMPI 2.0. (A) Mutations partitioned according to their origin, the number of altered residues, location within the complex, by the availability of additional kinetic and thermodynamic data, according to the experimental method used, and by category. (B) Distribution of ΔΔ*G*. (C) Source and target amino acids for single point mutations.

**Fig. 2.**
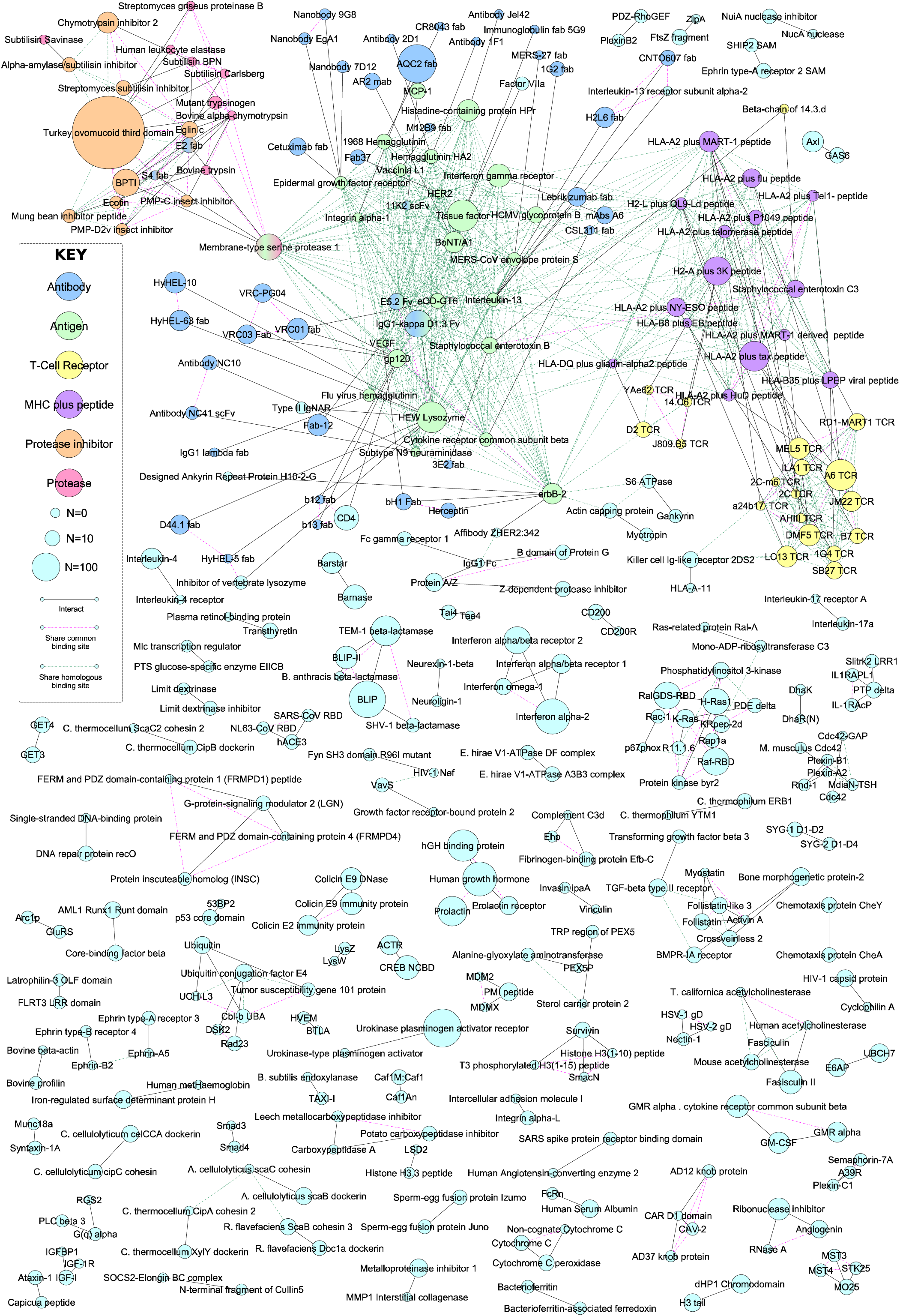
Overview of the interactions in SKEMPI. Nodes indicate proteins, scaled by the number of mutations of that protein and coloured according to category. Edges show direct interactions, as well as relationships between proteins that share a common or homologous binding site.

The entries also vary in the degree of structural order. While most correspond to interactions between folded domains, the database contains entries in which structuring occurs upon binding, such as protein-peptide interactions and, in the extreme case, the ACTR/NCBD interaction in which both binding partners become ordered upon binding [32]. Indeed, the requirements of having a structure in order to be included in the data set, *ipso facto* biases the data, and means that there is no representation of “fuzzy” complexes in which a diffuse structural ensemble in the bound state prevents the formation of a resolvable crystal.

Another source of variation is the origins of the interacting proteins. While entries range from viral and bacterial to the higher eukaryotes, biases are evident in the over-representation of model organisms including humans. With the exception of the pMHC-TCR, antibody-antigen and protease-inhibitor classes, most of the interactions are endogenous. Nevertheless, the set also includes exogenous interactions ranging from those between proteins from different individuals within the same species, namely the sex fusion proteins Juno and Izumo1 from human sperm and egg respectively [4], to host-pathogen interactions such as adenovirus and coronavirus interactions with human receptors during viral entry [65], [27], to the inhibition of acetyl-cholinesterase by the snake venom neurotoxin fasciculin [2]. For the pMHC-TCR interactions there are a variety of presented antigens, including exogenous viral peptides and gluten, as well as endogenous autoimmune and cancer peptides. The antibody interactions include pathogen antibodies, as well as antibodies raised and optimised to target extracellular therapeutic targets. The protease interactions are mostly exogenous, arising from their inherent cross-reactivity due to the convergent evolution of their canonical inhibitory loop.

### B. Notable studies comprising SKEMPI 2.0

The investigations from which SKEMPI data is derived are diverse, spanning many biological processes and reported in 295 publications including systematic scans, alanine and homolog scanning, design studies including computational design and designs derived from phage display, double mutant cycle studies, antibody engineering, biologic drug design and the evaluation of pathological mutations. The largest contribution comes from the group of the late Michael Laskowski Jr., a systematic study of all possible mutations at selected sites in the turkey ovomucoid third domain and its inhibitory interactions with four proteases [43], as well as studies of interactions of the same domain in other bird species and the design of ultra-high affinity broad-spectrum inhibitors. Substantial data also come from investigations into the inhibitory interactions of class A *β*-lactamases from the groups of Gideon Schreiber (e.g. [62]) and Timothy Palzkill (e.g. [10]), as well as cytokine receptor interactions, in particular studies of type I interferons also from the Schreiber group (e.g. [63]) and that of K. Christopher Garcia [70], but also the study of the GM-CSF / GMR*α* interaction from the group of Michael W. Parker [9]. Other prominent sources of data are studies into hormone receptor interactions, in particular the human growth hormone receptor from the group of Jim Wells (e.g. [14]) and the prolactin receptor from the group of Michael E. Hodsdon [35], as well as studies into antigen recognition including the combined computational and experimental design study to enhance affinity of the AQC2 antibody to integrin *α*-1 from the group of Herman Van Vlijmen [12], the dissection of the interactions of broadly neutralising antibodies targeting HIV gp120 [11] from the group of Richard A. Friesner, and various investigations from the group of Roy A. Mariuzza (e.g. [15]). Also notable is the alanine scanning of the urokinase-type plasminogen activator and its receptor from the group of Michael Ploug [21], studies of Ras effector interactions from the group of Christian Herrmann (e.g. [34]), investigations of pMHC/TCR interactions from the group of Brian Baker (e.g. [59]), and investigations into the congate and non-cognate recognition of *E. coli* colicin DNase bacteriotoxins by their immunity proteins from the group of Colin Kleanthous (e.g. [40]).

### C. Range and error

#### Range

The changes in binding free energy upon mutation range from −12.4 to 12.4 kcal.mol^−1^, as in SKEMPI 1.1, with Δlog_10_ *k*_on_ ranging from −3.6 to 2.4, Δlog_10_ *k*_off_ ranging from −6.0 to 6.8, ΔΔ*H* ranging from −18.3 to 26.5 kcal.mol^−1^, and ΔΔ*S* ranging from −61 to 80 cal.mol^−1^.K^−1^. Around 60 mutant are very destabilising, reducing binding energy by 8 kcal.mol^−1^ or more. These are all in enzyme/inhibitor complexes such as the inhibition of acetylcholinesterase by the snake venom fasciculin, or the inhibition of enzymes which would be detrimental should they unbind and become active in the wrong location, such as nucleases (barnase / barstar, colicin E9 DNase / Im9, RNase A / angiogenin) and proteases (such as trypsin / BPTI). These interactions tend to be around picomolar affinity and are at the upper limit of what can be detected, due to the time required to reach equilibrium and the low concentrations required by the mass action law to probe informative regions of the binding curve. These very destabilising mutations reduce affinity into the micromolar range, near the lower limit of what can be quantified using standard methods. As a consequence of both mutant and wild-type affinities being near detection thresholds, errors in these entries are typically large. Further, while some mutations may cause changes in affinity larger than seven orders of magnitude, the absence of affinities for such mutations in the benchmark can be explained by the fact that such mutations would involve affinities beyond the upper or lower limit. Indeed, there are new entries in which single or double substitutions reduce binding from tens of picomolar to having no detectable binding. For many of the highly destabilising mutations a crystal structure for the mutant has also been solved, and the 30 most stabilising mutations in the database (ΔΔ*G* ¡ −5 kcal.mol^−1^) consist of the reverse mutation applied to these structures. These are mostly single or double mutants, but include mutations to up to 27 residues of the non-cognate Colicin E2/Im9 complex, which move it toward the cognate E9/Im9 in sequence space [40].

#### Errors

Standard errors in *K_D_* are typically reported in the order of 50%, around 0.25 kcal.mol^−1^. These estimates are derived by repeat measurements using the same equipment, environment and protocol, and thus do not include errors arising from systematic bias. Such biases can, however, be estimated from pairs of entries in which the same mutation is evaluated by different groups or using different techniques. For 84% of 1741 such pairs, both entries give a ΔΔ*G* value within 1 kcal.mol^−1^ of each other. For 704 pairs for which kon is available for both, 80% have Δlog_10_ (*k*_on_) within 0.5 of each other. For 702 pairs for which ko _ff_ is available, 83% have Δlog_10_ (*k*_off_) within 0.5. For the 62 pair with both ΔΔ*H* and ΔΔ*S* values, 61% have ΔΔ*H* within 3.0 kcal.mol^−1^ of each other and 58% have ΔΔ*S* within 10 cal.mol^−1^.K^−1^ of each other.

### D. Mutant Cycles

Within SKEMPI some entries can be combined to construct mutant cycles, which quantify the interactions between residues, the dependence of these interactions on other residues, and other higher order effects. The most common instances are double mutant cycles, where affinities are available for the wild type, A, B and AB mutations, of which there are 610 examples. Of these, 53 involve at least one mutant for which binding was not observed, or only an inequality is available, and 235 involve mutations reported in the same reference, and thus the affinities are likely to have been measured using the same technique and conditions. A further 218 double mutant cycles can be constructed in the background of a third mutation (i.e. C, AC, BC and ABC mutations are available), of which 209 are not composed of non-binding mutations or mutations with inequalities, and 131 involve affinities coming from the same reference. Of the 766 double mutant cycles containing neither inequalities nor non-binding mutants, a number of parameters can be calculated, including ΔΔ*G_ab_*_→_ *_Ab_*, ΔΔ*G_ab→aB_* and ΔΔ*G_ab_*_→_ *_AB_*, the binding free energy change of both single and the double mutation respectively, as well as ΔΔ*G_aB_*_→_ *_AB_* and ΔΔ*G_Ab_*_→_ *_AB_*, the energy of a single mutation within the context of the other mutation, and ΔΔ*G_int_* = ΔΔ*G_aB_*_→_ *_AB_* − ΔΔ*G_ab_*_→_ *_Ab_* = ΔΔ*G_Ab_*_→_ *_AB_* − ΔΔ*G_ab_*_→_ *_aB_*, the interaction energy of the two mutations [26]. From these, it can be deduced that 345 are additive (ΔΔ*G_int_* < 0.5kcal.mol^−1^). Of the non-additive cycles, 293 exhibit tighter binding in the double mutant than the sum of the single mutants (positive epistasis), of which 6 result in even tighter binding than individual effects of two single mutations that strengthen the interaction (synergistic positive), while 273 correspond to double mutants which reduce binding by less than the sum of two single mutants which reduce binding (antagonistic positive). Similarly, 128 cycles have double mutants exhibiting weaker binding than the sum of the two single mutants (negative epistasis), of which 58 contain two destabilising single mutations (synergistic negative) and 26 contain two stabilising single mutations (antagonistic negative). The range of ΔΔ*G_int_* values rarely fall outside of the −5 to 3 kcal.mol^−1^ range. Of the 421 non-additive cycles, 151 show noticeable sign epistasis, in which the sign of the effect of either the A or B mutation flips depending on the presence or absence of the background mutation (i.e., for the A mutation, |ΔΔ*G_ab_*_→*Ab*_| > 0.2 kcal.mol^−1^ and |ΔΔ*G_aB_*_→*AB*_| > 0.2 kcal.mol^−1^ and |ΔΔ*G_ab_*_→*Ab*_ − ΔΔ*G_aB_*_→*AB*_| > 0.4 kcal.mol^−–1^). Of these, 38 correspond to mutations which destabilise the complex in the presence of the background mutation, but stabilise in its absence (destabilising sign epistasis), while 113 correspond to mutations which stabilise the complex in the mutant background but otherwise destabilise the complex (stabilising sign epistasis). Only 8 cycles exhibit the more extreme reciprocal sign epistasis, which in 6 cases are where both single mutations are stabilising (< –0.2 kcal.mol^−1^), but the double mutant is destabilising (> 0.2 kcal.mol^−1^), and the remaining two correspond to two destabilising mutations (> 0.2 kcal.mol^−1^) for which the double mutation is stabilising (< –0.2 kcal.mol^−1^). The types of substitutions that can give rise to extreme effects such as stabilising reciprocal sign epistasis can be illustrated with the Mlc-IIB^Glc^ interaction in *E. coli* [52], in which the removal of the F136 side-chain of MlC creates a large cavity at the binding interface, the addition of a phenylalanine at the A451 position of IIBG^l^c creates a large clash, however the double mutation creates an interaction that is even more stable than the wild-type by creating an anchor residue across the binding interface in which the cavity in MlC is filled by the new side-chain of IIB.

Higher order interaction terms can be garnered from higher cycles, such as triple mutant cubes, constructed from the energies of the wild-type, three single mutants, three corresponding double mutants and the triple mutant [26]. In SKEMPI, 45 triple mutant cubes can be made, with 10 coming from the same reference. For these, third order interaction energies fall within the −1 to 1 kcal.mol^−1^ range. For fourth order interactions, constructed from energies of the wild-type, four single mutants, six double mutants, four triple mutants and the quadruple mutant, 14 examples exist within SKEMPI. However, care should be taken in ascribing meaning to fourth order residue coupling energies due to the accumulations of errors, which in these cases are exacerbated by the affinities having been reported in different publications. No fifth or higher order interactions are present.

### E. The SKEMPI website

The database is accessible online at https://life.bsc.es/pid/skempi2/, where the raw CSV (comma-separated values) file containing all the data can be downloaded. The data can also be browsed online, ordered and searched by any field, such as the experimental method or the location of the mutation, or searching for a specific protein by its name or PDB code, and structures may be visualised. Other pages on the web site offer a summary of the data, an FAQ and help page, and a page for user contributions which will be evaluated to appear in future releases. Before you begin to format your paper, first write and save the content as a separate text file. Keep your text and graphic files separate until after the text has been formatted and styled. Do not use hard tabs, and limit use of hard returns to only one return at the end of a paragraph. Do not add any kind of pagination anywhere in the paper. Do not number text heads-the template will do that for you.

Finally, complete content and organizational editing before formatting. Please take note of the following items when proofreading spelling and grammar:

### F. Abbreviations and Acronyms

Define abbreviations and acronyms the first time they are used in the text, even after they have been defined in the abstract. Abbreviations such as IEEE, SI, MKS, CGS, sc, dc, and rms do not have to be defined. Do not use abbreviations in the title or heads unless they are unavoidable.

